# CRISPRi-TnSeq: A genome-wide high-throughput tool for bacterial essential-nonessential genetic interaction mapping

**DOI:** 10.1101/2023.05.31.543074

**Authors:** Bimal Jana, Xue Liu, Julien Dénéréaz, Hongshik Park, Dmitry Leshchiner, Bruce Liu, Clément Gallay, Jan-Willem Veening, Tim van Opijnen

## Abstract

Genetic interaction networks can help identify functional connections between genes and pathways, which can be leveraged to establish (new) gene function, drug targets, and fill pathway gaps. Since there is no optimal tool that can map genetic interactions across many different bacterial strains and species, we develop CRISPRi-TnSeq, a genome-wide tool that maps genetic interactions between essential genes and nonessential genes through the knockdown of a targeted essential gene (CRISPRi) and the simultaneous knockout of individual nonessential genes (Tn-Seq). CRISPRi-TnSeq thereby identifies, on a genome-wide scale, synthetic and suppressor-type relationships between essential and nonessential genes, enabling the construction of essential-nonessential genetic interaction networks. To develop and optimize CRISPRi-TnSeq, CRISPRi strains were obtained for 13 essential genes in *Streptococcus pneumoniae,* involved in different biological processes including metabolism, DNA replication, transcription, cell division and cell envelope synthesis. Transposon-mutant libraries were constructed in each strain enabling screening of ∼24,000 gene-gene pairs, which led to the identification of 1,334 genetic interactions, including 754 negative and 580 positive genetic interactions. Through extensive network analyses and validation experiments we identify a set of 17 pleiotropic genes, of which a subset tentatively functions as genetic capacitors, dampening phenotypic outcomes and protecting against perturbations. Furthermore, we focus on the relationships between cell wall synthesis, integrity and cell division and highlight: 1) how essential gene knockdown can be compensated by rerouting flux through nonessential genes in a pathway; 2) the existence of a delicate balance between Z-ring formation and localization, and septal and peripheral peptidoglycan (PG) synthesis to successfully accomplish cell division; 3) the control of c-di-AMP over intracellular K^+^ and turgor, and thereby modulation of the cell wall synthesis machinery; 4) the dynamic nature of cell wall protein CozEb and its effect on PG synthesis, cell shape morphology and envelope integrity; 5) functional dependency between chromosome decatenation and segregation, and the critical link with cell division, and cell wall synthesis. Overall, we show that CRISPRi-TnSeq uncovers genetic interactions between closely functionally linked genes and pathways, as well as disparate genes and pathways, highlighting pathway dependencies and valuable leads for gene function. Importantly, since both CRISPRi and Tn-Seq are widely used tools, CRISPRi-TnSeq should be relatively easy to implement to construct genetic interaction networks across many different microbial strains and species.

## Introduction

Most bacterial genomes contain many genetic elements with unknown function. Uncovering which genetic elements in a genome work together (i.e., interact) and how they are involved in generating a phenotype is thereby a key challenge. However, the rate at which genome-wide phenotyping/genomics tools are associating genes with a (new) function is not keeping pace with the rate at which new genetic material is sequenced. A variety of Transposon Insertion Sequencing tools including Transposon Insertion Sequencing (Tn-Seq)^1^ and more recently CRISPR interference (CRISPRi)^2^ have been used in different bacterial strains and species to associate genetic elements with a specific environmental and/or genetic perturbation, which can in turn be used as a lead for gene function. For instance, transposon and CRISPRi libraries grown in the presence of different antibiotics or host-like conditions have been used to identify genes associated with antibiotic transport, cell wall synthesis, membrane integrity, regulation, and global responses to overcome stress^3–5^. Both tools have their advantages, for instance Tn-Seq is often more easily implementable than CRISPRi, while the latter tool is able to target essential genes that remain out of reach of Tn-Seq. Importantly, they both lack the ability to easily map genetic-interactions on a genome-wide scale, which can make it hard to understand how genetic elements fit within the larger genomic context and thereby untangle their function.

One of the most comprehensive genetic interaction networks has to date been constructed for *Saccharomyces cerevisiae* through synthetic genetic array (SGA) analyses^6^. SGA is a tool that allows for sampling of double gene knockouts, and temperature-sensitive (TS) mutants^7^ on a genome-wide scale. Yeast networks generated with SGA have been used to develop hierarchical genetic models of cellular functions, comprising small to large functional modules; protein complexes, pathways, biological processes, and cellular compartments^8^. Additionally, through ‘guilty by association’ analyses, in which genes with unknown function interact with genes with a known function and are thereby hypothesized to share a similar function, new leads for gene function have been uncovered, including for genes important for critical processes such as cell division, DNA replication and transcription^8^. Genetic interaction networks have been generated for human K562 and Jurkat cells by knocking down genes in combination using dual-CRISPRi, which highlights the interdependency between core pathways like cholesterol biosynthesis and DNA repair and identified potential targets for anti-cancer combination therapy^9^. Moreover, synthetic genetic array (eSGA) technology in *E. coli,* which through conjugation between gene deletion and/or hypomorph strains enables double mutant generation on a genome-wide scale, has been used to map genetic interaction networks that have helped untangle different pathways including those involved in synthesizing iron-sulfur (Fe-S) clusters^10^. While these array-based approaches are powerful and highlight the importance of generating interaction networks for phenotyping analyses, they are only available for a select number of model strains and organisms^11, 12^, and they lack the ability to easily scale across many bacterial strains and species.

Transposon insertion sequencing-based methods, including Tn-Seq, have enabled identification of essential genes in non-model microorganisms, as well as studying the contribution of individual genes to phenotypes on a genome-wide scale^1, 3, 13–15^. Additionally, Tn-Seq has enabled genetic interaction mapping by constructing transposon mutant libraries in the background of a deletion mutant of interest. This has led to the identification of a variety of new functions including regulatory relationships among genes^3, 13^, pathway and transporter redundancies^16^, membrane integrity genes^17^ and genes involved in cholesterol processing^18, 19^. However, although essential genes are hubs within genetic interaction networks connecting critical genes and processes^8^, a key limitation of the deletion array or transposon insertion-based approaches is that they cannot sample essential genes, as they are inviable as knockouts^20^. To, at least partially, bypass this issue, chemical-genetics approaches have been used by targeting essential genes with antimicrobial compounds while simultaneously knocking out nonessential genes, enabling mapping of genetic interactions with essential genes^21^. However, this approach only goes so far, as compounds are available against only a limited number of essential gene products, and they often trigger off-target effects therefore lacking specificity. The development of CRISPRi has made studying essential gene function in bacteria possible^22–24^ by using the complex of a target gene specific single <u>g</u>uide RNA and a catalytically dead Cas protein to block transcription elongation of a target gene, resulting in (potentially) tunable gene knockdown. CRISPRi thereby enables gene knockdown of both nonessential and essential genes, which for instance has led to uncovering new genotype-phenotype relationships for genes involved in cell wall synthesis, division, and competence^25–27^.

While CRISPRi often needs extensive development to implement it in a new strain or species, and Tn-Seq cannot sample essential genes, we here exploit the advantages of either approach by adding the new approach CRISPRi-TnSeq to the genomics toolbox. Specifically, by taking advantage of the easily implementable tool Tn-Seq and combining it with CRISPRi, genetic interaction mapping is enabled between nonessential and essential genes. As a proof of principle, CRISPRi-TnSeq is implemented in *Streptococcus pneumoniae* to map genome-wide interactions for 13 essential genes by screening ∼24,000 gene-gene pairs, which leads to 1334 significant essential-nonessential genetic interactions. By performing extensive analyses and validation experiments, including chemical genetics and detailed knockout and microscopy studies we show the applicability of CRISPRi-TnSeq in constructing genetic interaction networks. These data are subsequently mined to highlight new biology, including the identification of a set of pleiotropic genes that provide robustness to the organism, the ability of nonessential genes to compensate for essential genes, the identification of new roles and relationships for genes involved in cell wall synthesis and division, the importance of cyclic-di-AMP in controlling turgor and thereby cell wall synthesis, and a functional link between DNA replication and cell division. Lastly, we provide a clear experimental and analytical roadmap to implement CRISPRi-TnSeq across many other bacterial species with the aim to accelerate genotype-phenotype discovery.

## Results

### Combining CRISPRi and Tn-Seq to map essential-nonessential gene interaction networks

CRISPRi-TnSeq identifies genetic interactions between essential genes and nonessential genes through the knockdown of a targeted essential gene (CRISPRi) and the simultaneous knockout of a nonessential gene (Tn-Seq). CRISPRi-TnSeq thereby identifies, on a genome-wide scale, synthetic and suppressor-type relationships between essential and nonessential genes, enabling the construction of essential-nonessential genetic interaction networks. CRISPRi-TnSeq follows 5 main steps: **1**) A bacterial strain (here we used *S*. *pneumoniae* D39) is engineered to contain *dcas9* on the chromosome under control of the IPTG inducible P*lac* promoter. In the same genome a constitutive P3 promoter drives the expression of a specific single guide (sg) RNA targeting an essential gene. Induction of dCas9 with IPTG thereby allows conditional and tunable knockdown of a target gene^26^ (Fig. 1.1); **2)** In the background of such a CRISPRi strain a transposon mutant library is introduced using the Magellan6 transposon as described previously^13^ (Fig. 1.2); **3)** Individual libraries are grown in a condition of interest and in the presence and absence of a sub-inhibitory concentration of IPTG (Fig. 1.3); **4)** Tn-Seq sample preparation, sequencing and analyses are performed to calculate the fitness effect of each nonessential gene knockout in the genome with and without knockdown of the targeted essential gene (Fig. 1.4); **5)** Negative and positive interactions are scored as a significant deviation from the expected multiplicative of the fitness of the knocked-out nonessential and knocked-down essential gene, enabling the mapping of a genetic interaction network (Fig. 1.5).

**Figure 1.**
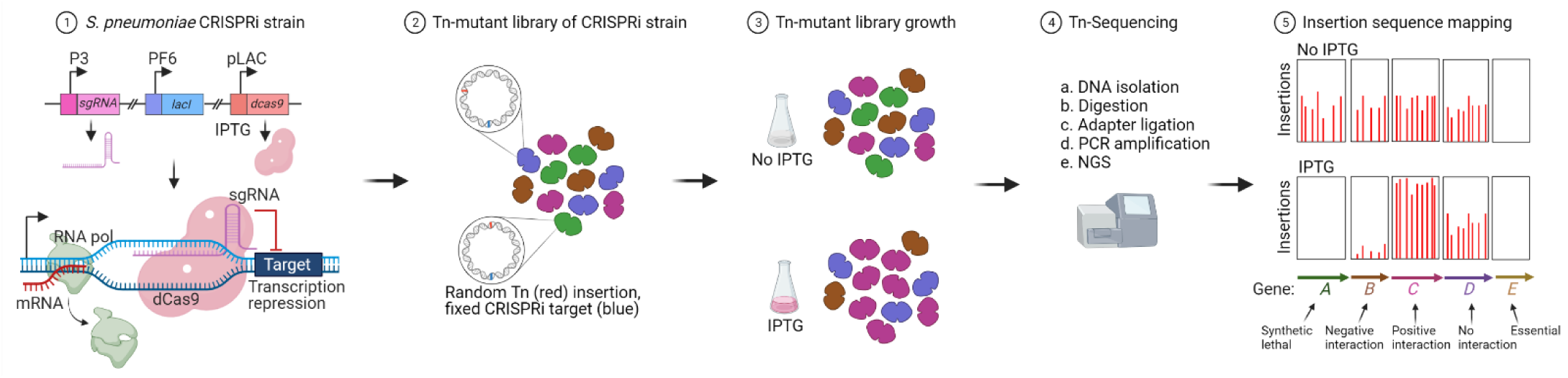
Schematic overview of CRISPRi-TnSeq. **1.** The *S. pneumoniae* D39 CRISPRi strain carries a target gene specific sgRNA under a P3 constitutive promoter, *dcas9* under the P*lac* inducible promoter, which is blocked by LacI expressed from the constitutive promoter PF6. Addition of IPTG to the medium triggers expression of dCas9, which leads to conditional knockdown of the targeted essential gene. **2.** Transposon mutant libraries are introduced in a CRISPRi essential-gene knockdown strain, creating genome-wide non-essential gene knockout libraries in the background of a strain in which an essential gene can be knocked down through the addition of IPTG. **3.** A transposon mutant library is grown for multiple generations with and without IPTG. **4.** DNA is isolated from each population before and after growth in the presence and absence of IPTG, followed by TnSeq analysis. **5.** Transposon insertion sites are mapped to the genome and the fitness effect for each non-essential gene in the presence and absence of IPTG is calculated, leading to phenotypes including synthetic, negative, positive or no interaction.

### Essential gene knockdown and growth inhibition are stable in transposon mutant libraries

To develop and optimize CRISPRi-TnSeq, we selected a set of 13 essential genes, involved in different biological processes including metabolism, DNA replication, transcription, cell division and cell envelope synthesis, and obtained a CRISPRi strain for each of them (Fig. 2A, Supplemental Table 1). Transposon mutant libraries consisting of ∼60,000 mutants were constructed in each individual CRISPRi strain. To confirm that gene knockdown is not affected by the introduction of a transposon library in a CRISPRi strain’s background, six libraries were grown in the absence and presence of sub-inhibitory concentration(s) of IPTG to establish growth curves and quantitative PCR was performed to confirm target gene knockdown. Increased growth inhibition is observed with increased concentrations of IPTG for essential genes, except *clpP* which is not essential for *in vitro* growth of *S. pneumoniae* D39 (Fig. 2B). Additionally, gene expression of CRISPRi targeted genes is significantly reduced with the addition of IPTG and is titratable for most genes with a notable exception for *parC* (Fig. 2B, Supplemental Table 2). This means that while at 100 mM IPTG *parC* gene-expression can be completely repressed, the effect on growth is much less dramatic and abrupt. This indicates that the effect of essential gene knockdown on growth can be target specific and does not necessarily have an immediate impact, which has similarly been shown for DNA replication associated genes in *Mycobacterium tuberculosis*^28^. Overall, these data show that partial knockdown of essential genes can be achieved efficiently in the background of a transposon mutant library.

**Figure 2.**
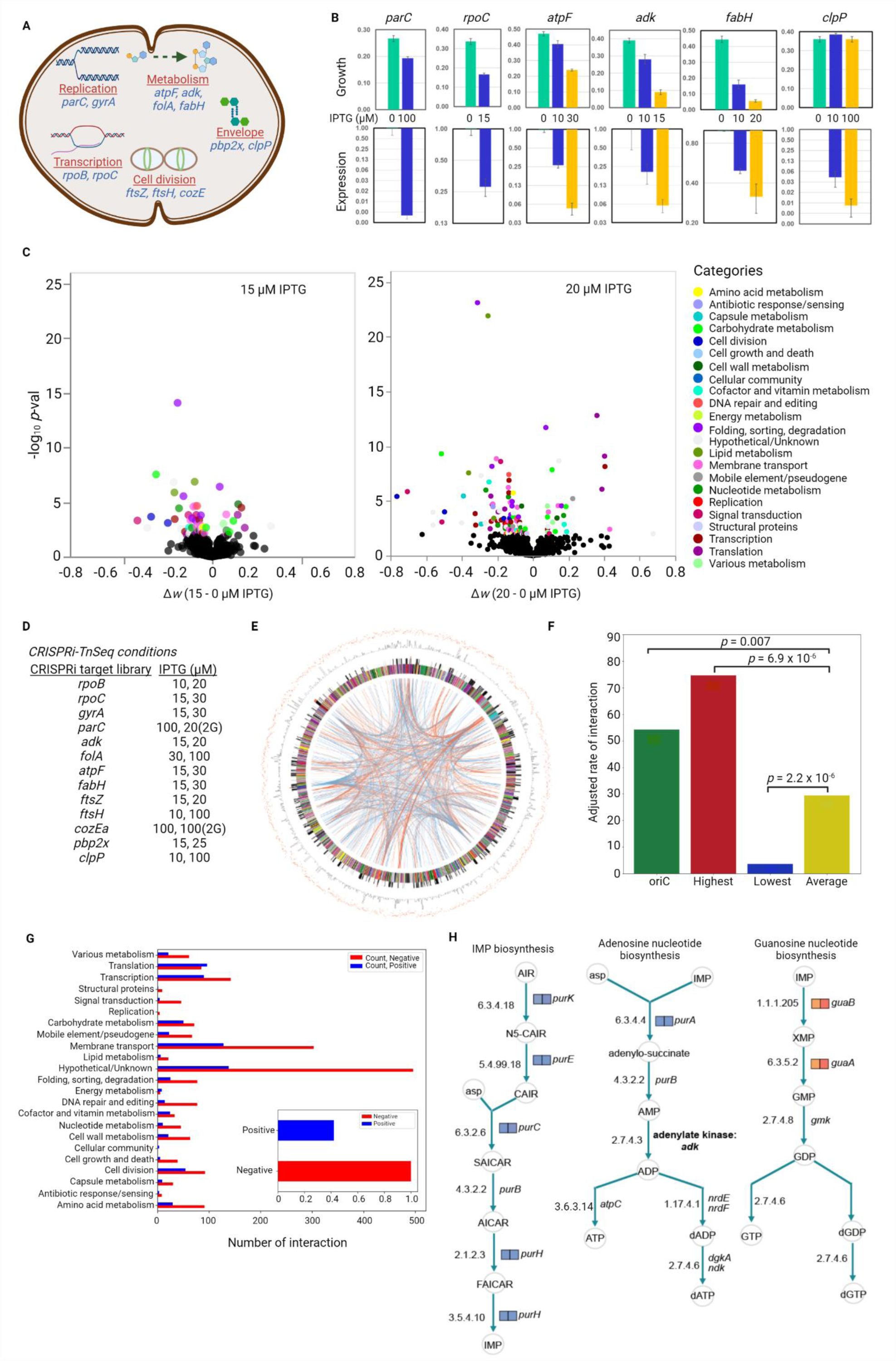
CRISPRi-TnSeq identifies essential-nonessential genetic interactions at genome-scale. **A**. Transposon mutant libraries were constructed in 13 *S. pneumoniae* D39 CRISPRi strains, targeting essential genes involved in different bioprocesses. **B**. Sub-inhibitory IPTG concentrations inhibit growth (upper panel) and knockdown (lower panel) gene expression, measured by RT-qPCR for libraries of 6 CRISPRi strains in C+Y medium, validating functionality of the CRISPRi system in Tn-mutants. **C**. Example volcano plots for *fabH* CRISPRi strain. The transposon mutant library of the strain was grown at 0, 15 and 20 μM IPTG, followed by CRISPRi-TnSeq analysis. Increased knockdown due to increased IPTG highlights the identification of an increased number of genetic interactions (colored by bioprocesses). **D**. List of transposon mutant libraries/CRISPRi strains and the corresponding IPTG concentrations used in the CRISPRi-TnSeq screen. A library was selected sequentially in two rounds of growth (2G) in the presence of IPTG when CRISPRi knockdown impacted growth minimally. **E**. Genome-wide interactions between essential genes and non-essential genes are depicted in a circos-plot. The red and blue lines represent positive and negative interactions, respectively. The figure highlights the distribution of interactions across the entire genome. A larger version of the figure, highlighting additional information is shown in Supplemental Figure 3. **F**. Some areas of the genome carry significantly higher or lower numbers of interactions compared to the average number of interactions. Interactions per 50 kb and the *p*-val compared with the average interactions are shown. **G**. Genetic interactions are distributed over different biological processes. Negative interactions (red) are more prevalent then positive (blue) interactions. **H**. Purine biosynthesis pathways become important when *adk* is knocked down. Nonessential genes involved in AMP biosynthesis are identified with negative genetic interactions in the presence of 15 or 20 µM IPTG presented by blue boxes. In contrast, nonessential genes involved in GMP biosynthesis are disadvantageous under *adk* knockdown therefore identified with positive genetic interaction presented by red boxes.

### CRISPRi-TnSeq identifies nonessential genes mechanistically linked to essential genes

To test the ability of CRISPRi-TnSeq to identify genetic interactions, transposon mutant libraries were grown in the absence and presence of two different IPTG concentrations. Subsequently, Tn-Seq analysis was performed^13, 14^ resulting in a fitness value (i.e., growth rate) for each nonessential gene in the genome. Importantly, in the absence of IPTG fitness represents the growth rate solely affected by gene-disruption because of the transposon insertion, while in the presence of IPTG the effect on fitness is a combination of the nonessential gene-disruption and the essential gene-knockdown. Genetic interactions are subsequently defined as negative if fitness in the presence of IPTG (*W_IPTG_*) is significantly smaller than fitness in the absence of IPTG (*W*_noIPTG_) and thereby deviates from the expected multiplicative fitness (i.e., the expected fitness from combining the individual effects of the essential gene knockdown and the nonessential gene disruption), while a positive interaction is defined as *W_IPTG_* being significantly higher than *W*_noIPTG_. Using the 13 CRISPRi strains (Fig. 2D) we screened ∼24,000 gene-gene pairs and identified 1,334 significant genetic interactions (Supplemental Fig. 1, Supplemental Table 3), which could be divided into 754 negative and 580 positive genetic interactions. Permutation testing indicates that the experimental dataset has a non-random distribution (i.e., a significantly higher number of negative genetic interactions are present in the experimental datasets (Supplemental Fig. 2)), which resembles a distribution of genetic interactions observed in synthetic genetic array screens of *Escherichia coli*^10^.

We have previously shown that a fitness effect of a genetic insult (e.g., a gene disruption) can become more pronounced with increasing stress, for instance a gene important for overcoming antibiotics will display increasing importance (e.g., decreasing fitness) with increasing amounts of antibiotics^4^. Measuring fitness under different stress levels can thus determine the fitness importance and reproducibility of a gene or genetic interaction. Here, experiments were performed at two sub-inhibitory IPTG concentrations, where it is expected that interactions identified at the lowest concentration should be mostly reproducible at the higher concentration (i.e., increased essential gene knockdown leads to increased stress), while simultaneously identifying additional interactions. Overall, reproducibility was high with an average of 65% overlap between IPTG concentrations (Supplemental Table 4), with the highest overlap for *fabH*; i.e., out of 51 interactions identified at 15 mM, 42 were also identified at 20 mM (82% overlap), while simultaneously identifying 82 additional interactions at 20 mM (Supplemental Table 4).

All genetic interactions were mapped on a circular genome and can be found across all gene categories (Fig. 2E). Nonessential genes located around *oriC* possess a significantly higher number of interactions (Fig. 2F, Supplemental Fig. 3), which is likely due to increased gene dosage around *oriC* to overcome replication stress^29^. Furthermore a 50 kb region spanning genes SPD2014-SPD2064 has the highest number of interactions (Fig. 2F, Supplemental Table 5), consisting of genes involved in stress response, cell division, competence, and regulation. Negative interactions dominate positive interactions in all categories except for translation (Fig. 2G). This suggests that shutting down one of the most energy costly systems can help cells in certain situations to cope better with stress experienced from essential gene knockdown, especially those involved in replication, transcription, and energy metabolism. Furthermore, genetic interactions with essential gene *atpF,* part of the ATP synthase complex, are mostly positive (Supplemental Table 3). We and others have shown that a reduction in ATP can trigger tolerance towards antibiotics including DNA and protein synthesis inhibitors^4^. Reducing ATP may thus also buffer against more general stress, in this case stress coming from knocking out genes involved in translation, carbohydrate or nucleotide metabolism. Moreover, as ATP synthase in *S. pneumoniae* is used to maintain pH homeostasis, *atpF* knockdown might increase intracellular pH and reduce growth^30^, which may additionally contribute to buffer against stress.

Gene set enrichment analysis shows that knockdown of an essential gene often leads to genetic interactions with functionally linked genes (Supplemental Fig. 4, Supplemental Table 6). For example, DNA repair and lipid metabolism genes are enriched in the *parC* and *fabH* knockdown datasets, respectively. Moreover, these functional connections are also present when individual genes are considered at the pathway level. For example, nonessential genes in the AMP biosynthesis pathway have a negative interaction with *adk*, while genes involved in GMP biosynthesis have a positive interaction (Fig. 2H). Overall, genes involved in purine biosynthesis are advantageous when *adk* is knocked down, whereas genes involved in purine degradation are disadvantageous (Supplemental Fig. 5). Since AMP is the substrate of adenylate kinase Adk to produce ADP, reduction in the levels of AMP substrate likely impact the growth strongly when *adk* is already knocked down. Therefore, compensatory pathways which are involved in AMP biosynthesis become crucial. Overall, these results suggest that CRISPRi-TnSeq can identify nonessential genes that are mechanistically linked to corresponding essential genes.

### CRISPRi-TnSeq cross-referenced with antibiotic-TnSeq identifies functional modules

To further confirm that CRISPRi-TnSeq identifies biologically relevant interactions between essential and nonessential genes, we assessed the extent to which CRISPRi-TnSeq and antibiotic-TnSeq datasets overlap. While antibiotics often target essential gene products and/or processes, the ultimate cause of antibiotic mediated inhibition is often associated with the downstream impact on associated processes^31^. For instance, while fluoroquinolones like ciprofloxacin target DNA gyrase and/or topoisomerase to block DNA replication, inhibition of DNA repair can exacerbate the antibiotic’s effects due to its indirect ‘ability’ to increase DNA damage. We thus reasoned that the interactions between nonessential genes and a CRISPRi targeted essential gene, should overlap with interactions identified between nonessential genes and an antibiotic targeted essential gene product and/or process. We chose 8 antibiotics that target a specific essential gene product and/or process, including replication, transcription, cell division, carbohydrate, nucleotide, and cell wall metabolism (Fig. 3A). Similar to previous experiments^4^ transposon mutant libraries were grown in the presence of an antibiotic at a concentration that reduces overall growth by 30-50%, followed by Tn-Seq analysis. Genes in the same pathway, protein complex or biological process should, due to their shared function, have similar fitness profiles when measured across a set of conditions (e.g., different antibiotics) as well as share similar genetic interaction profiles. Profile similarity analysis thus provides an opportunity to validate the dataset, as clustered genes should represent functionally linked gene-sets and/or operons^8^. Both the CRISPRi-TnSeq and the antibiotic-TnSeq datasets were used in hierarchical clustering (Supplemental Table 2, 7). First, target specific CRISPRi-TnSeq and antibiotic-TnSeq datasets cluster closely together (Fig. 3B, Supplemental Table 8). For example, the *rpoC*-CRISPRi (*rpoC*i) dataset clusters tightly with the rifampicin-TnSeq (rifampicin-Tn) dataset, *pbp2x*i with penicillin-Tn, *fabH*i with cerulenin-Tn, *atpF*i with mefloquine-Tn and *parC*i with ciprofloxacin (Fig. 3B). Furthermore, hierarchical clustering also identifies operons and modules of nonessential genes involved in a specific process. This includes a module of class A PBPs involved in cell wall synthesis (*pbp1b*-*pbp2a*-*pbp1a*), the *Ami* oligopeptide operon (*amiA*-*amiD*-*amiC*-*amiE*-*amiF*), a potassium transporter *(trkA*-*trkH*), and protein synthesis genes *mnmE* and *mnmG*. Moreover *smc*-*scpA*-*scpB* form a cluster that, to our knowledge, has not been previously identified in *S. pneumoniae* although was expected as bacterial condensins consist of the SMC-ScpA-ScpB triad^32^. Interestingly, the homologous MukB-E-F complex of *E. coli* plays an important role in localizing and coordinating the function of topoisomerase IV^33–35^. SMC has been shown to play a role in chromosome segregation in *S. pneumoniae*^36^ and its clustering with ScpA-ScpB during fluoroquinolone treatment suggests that it may also play a role in coordinating topoisomerase IV. Clustering of fitness and genetic interaction data can thus be leveraged to verify known associations between genes as well as identify new ones.

**Figure 3.**
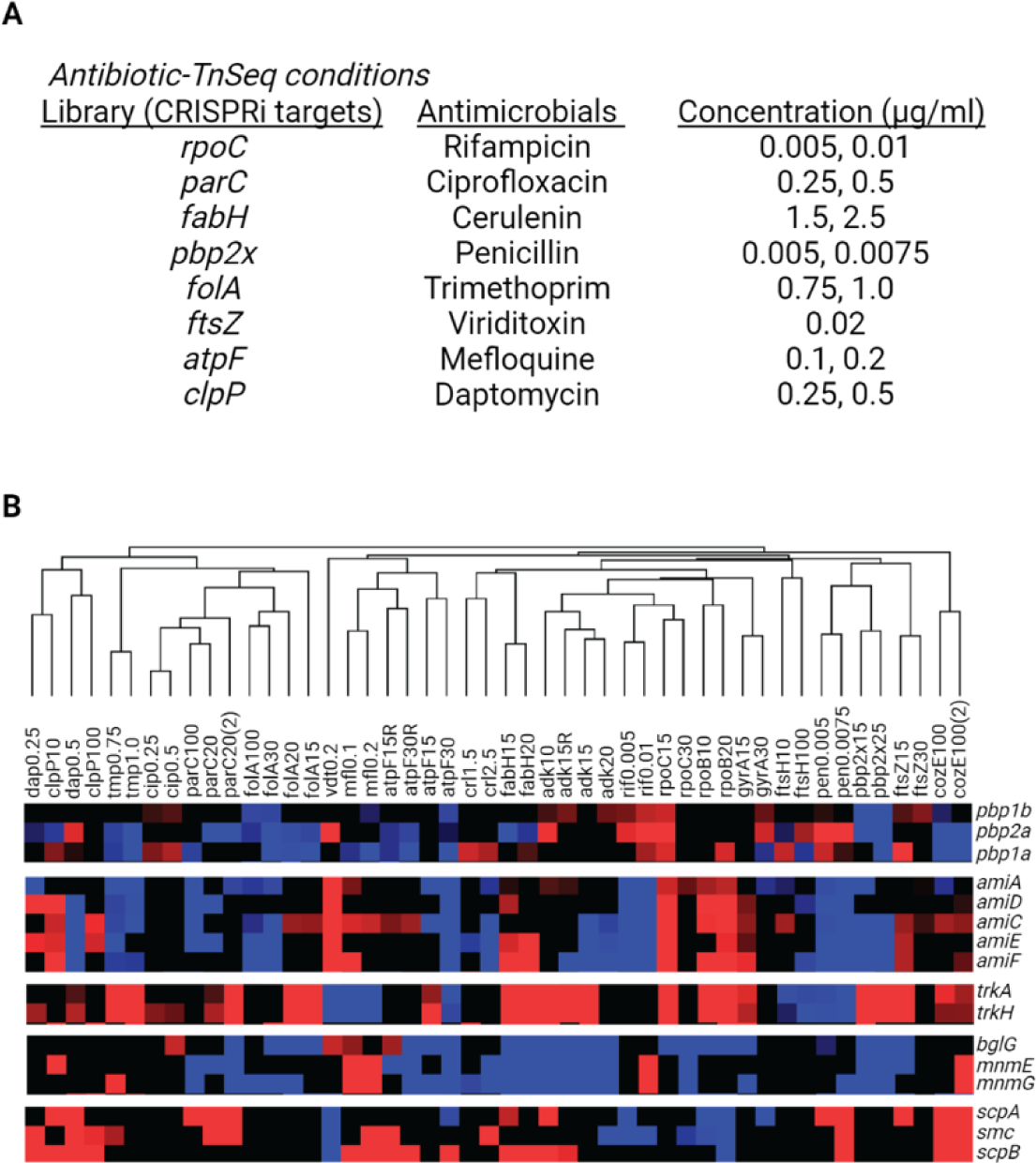
CRISPRi-TnSeq and antibiotic-TnSeq datasets cluster through overlapping targets. **A**. List of transposon mutant libraries of CRISPRi strains carrying a sgRNA targeting different essential genes (column 1), antibiotics that impair the function of the corresponding CRISPRi target (column 2), and the antibiotic concentrations used in antibiotic-TnSeq. **B**. Spearman’s rank correlation coefficient-based clustering of CRISPRi- and antibiotic-TnSeq datasets, identifies known and novel functional units of nonessential genes.

### *S. pneumoniae* contains pleiotropic genes that can modulate the experienced stress

Genetic interactions were mapped on a network containing 853 nonessential genes, which were ranked based on their total number of interactions (Fig. 4A). 17 of the highest ranked genes have interactions with more than half of the targeted essential genes (Fig. 4B). We validated 7 interactions (Fig. 4C, Supplemental Fig. 6) confirming that these genes interact with multiple essential genes and can thereby influence various bioprocesses. Seven of these genes are dominated by negative interactions, which includes critical regulators *ctsR* and *glnR*, the protease *clpC*, cell division gene *divIVA*, central nucleotide metabolism gene *purA*, phosphate transporter *phoU*, and *glnA*, which is central to glutamine metabolism and thereby the synthesis of amino acids and nucleotides. Knocking out these genes significantly increases the sensitivity of the organism to knockdown of essential genes from a wide variety of processes, making them the most pleotropic genes in the dataset. The ten remaining high degree genes have mostly positive interactions, and thus have a suppressor phenotype. Of this set, three are involved in protein synthesis (*mnmE*, *mnmG* and *rny*), three in transport (*psaA*, *psaC*, *gshY*), and two in metabolism (*ulaH*, *guaB*). When knocked out they can severely slow down growth and/or inhibit a specific process that may thereby mask the effect of essential gene knockdown, i.e., slowing down growth and/or a specific process can diminish the effects of experienced stress, which resembles the manner in which antibiotics can be tolerated by a bacterium by slowing down an antibiotic targeted/associated process^4^. These pleiotropic genes thereby tentatively function as genetic capacitors, which are genes that can dampen a phenotypic outcome, and introduce robustness into a biological system to protect against environmental or genetic insults. A canonical example of such a capacitor is heat shock protein 90 (HSP90) of *Saccharomyces cerevisiae*, a molecular chaperon that assists maturation of key regulatory proteins and thereby maintains many signaling networks and corresponding bioprocesses^37^. This network of pleotropic genes thus suggests *S. pneumoniae* contains a variety of nonessential genes that can work as genetic capacitors and assist in overcoming stress. Importantly, they could therefore also be considered as targets to sensitize the organism to various drugs. Note that BglG, an anti-terminator of operon *bglGFA* (SPD0501-3) which encodes enzymes involved in phosphorylation of sugar substrates, is also identified as having a suppressor phenotype. However, a *bglG* knockout is likely reducing the phosphorylation and transport of IPTG, the inducer of the CRISPRi system. It thereby reduces the activity of CRISPRi-mediated gene knockdown and should not be considered a suppressor.

**Figure 4.**
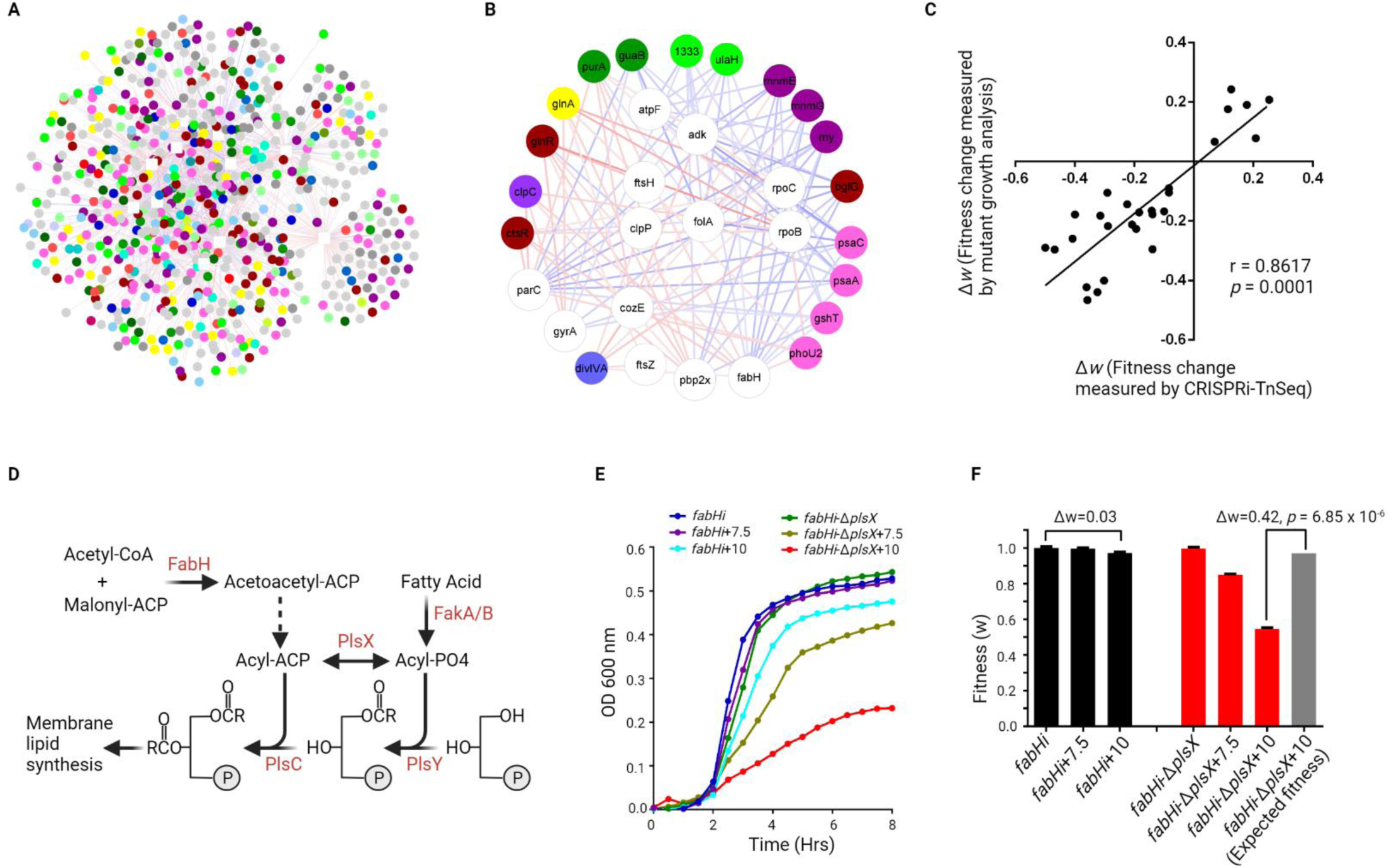
A CRISPRi-TnSeq essential-nonessential genetic interaction network. **A**. The genome-wide CRISPRi-TnSeq network contains 13 essential genes, 853 non-essential genes and 1334 genetic interactions. CRISPRi-target genes are presented as white squares, and nonessential genes as circles colored based on their bioprocess listed in Figure 2 with color code. Nodes are connected by colored edges which refer to negative (red) or positive (blue) interactions. **B**. A sub-network selected from the genome-wide network, highlights a set of 17 nonessential genes (colored circles) that ranked highest based on interactions with more than half of the targeted essential genes (white circles). These pleiotropic genes tentatively function as genetic capacitors and help protect against environmental or genetic insults. **C**. 27 genetic interactions were validated by constructing nonessential gene deletion mutants in the background of specific CRISPRi knockdown strains, followed by growth rate measurements. Measured growth rates were compared with the interactions of corresponding gene pairs identified in CRISPRi-TnSeq analysis, resulting in high reproducibility (Pearson correlation coefficient r = 0.86, *p* = 0.0001). **D**. Partial membrane lipid biosynthesis pathway showing the function and relationship of *plsX* and *fabH* and illustrating how *plsX* can partially compensate for *fabH* knockdown by increasing flux between Acyl-ACP and Acyl-PO4. **E**. Growth curves of *fabHi* with and without Δ*plsX.* While *fabHi* only has a slight growth defect in the presence of 10 µM IPTG, and there is no growth defect for Δ*plsX,* a significant growth defect occurs in *fabHi-*Δ*plsX* with 7.5 and 10 µM IPTG. **F**. Growth curves of *fabHi* and *fabHi-*Δ*plsX* transformed into a bar graph showing their observed relative fitness (*ΔW*) under each condition, as well as the expected multiplicative fitness of the double mutant. Comparisons between observed and expected relative fitness shows that the observed fitness is significantly lower than what would be expected under the multiplicative model (T-test P=6.85 x 10^-6^), indicating a strong negative interaction between both genes.

### CRISPRi-TnSeq essential-nonessential genetic interaction network validation

To further evaluate the quality of the CRISPRi-TnSeq dataset, 27 genetic interactions were validated by constructing nonessential gene deletion mutants in the background of specific CRISPRi knockdown strains (Supplemental Table 9). Growth rates were measured for each strain in the absence and presence of IPTG, thereby enabling calculations of each strain’s fitness affected by the nonessential gene-deletion (-IPTG), and simultaneous CRISPRi-mediated target knockdown (+IPTG) (Supplemental Table 10). Genetic interactions of essential and nonessential genes determined in individual growth studies correlated well with the interactions of corresponding gene pairs identified in CRISPRi-TnSeq analysis (r = 0.86, *p* = 0.0001; Fig. 4C, Supplemental Table 11). We further explored several of these interactions to obtain a better understanding of the interactions’ underlying biological mechanisms. We first explored a negative genetic interaction between *fabH*, an essential gene involved in initiating membrane lipid biosynthesis, and the nonessential gene *plsX,* which catalyzes the reversible formation between acyl-phosphate and acyl-ACP (Acyl Carrier Protein) (Fig. 4D). To validate this interaction, we constructed a Δ*plsX* mutant in a *fabH*-CRISPRi background (*fabH*i-Δ*plsX)* and determined growth of *fabH*i and *fabH*i-Δ*plsX* in the absence and presence of increased concentrations of IPTG (Fig. 4E). In the absence of IPTG, *fabH*i and *fabH*i-Δ*plsX* do not have a growth defect, confirming the Tn-Seq data that *plsX* by itself has no effect on growth under these conditions. At 7.5 and 10 mM IPTG, which leads to ∼2 to 3-fold reduction in expression of *fabH* (Fig. 2B) the effect on growth for *fabH*i is minimal. When screening for genetic interactions the null hypothesis is that most genes in a genome function independently and their combined fitness is that of the multiplicative of the individual effects. The multiplicative of combining *fabH*i with Δ*plsX* in the same background would thus predict that the effect on growth would mimic that of *fabH*i. However, growth of *fabH*i-Δ*plsX* at 7.5 and 10 mM IPTG is significantly inhibited compared to that of *fabH*i at the same IPTG knockdown levels (Fig. 4E). Converting growth curves to fitness further highlights the strong impact of *fabH* knockdown on the growth of Δ*plsX* (Fig. 4F). Growth analysis thus confirms the CRISPRi-TnSeq predicted negative genetic interaction between *fabH* and *plsX,* which stems from their ability to modulate cellular levels of Acyl-ACP, an essential intermediate in membrane lipid synthesis (Fig. 4D.). This means that when FabH is knocked down acyl-ACP levels are reduced which can be supplemented through PlsX’s ability to generate acyl-ACP through the reversible transfer of acyl-PO4 (Fig. 4D). Thus, while PlsX is dispensable, it can compensate for a loss in FabH functionality by diverting fatty acid and acyl-phosphate towards acyl-ACP production.

### Septal and Peripheral PG synthesis need to be balanced to retain proper cell shape and viability

Cell division is a complex process requiring the action of multiple pathways including cell wall (CW) synthesis. CW synthesis can be subdivided into peripheral and septal peptidoglycan (PG) synthesis. Peripheral PG synthesis enables cell elongation, important for building the progeny cell, and in which PBP2b and PBP1a are key nodes^38^. In contrast, PBP2x is a key node in septal PG synthesis that drives cell division^38^. How septal and peripheral PG synthesis at these two different cellular locations are orchestrated and whether they are dependent on each other/interact is unclear. By performing CRISPRi-TnSeq in a PBP2x CRISPRi knockdown strain we find that PG precursor synthesis genes *murM*, *murN*, and *murA1* become increasingly important, i.e., fitness significantly decreases, when PBP2x is knocked down and one of the precursor genes is knocked out (Fig. 5A). In contrast, when *pgdA* (peptidoglycan-N-acetylglucosamine deacetylase A) is knocked out, preventing deacetylation of PG, the negative effect of *pbp2x* knockdown, which results in slow growth, and elongated cells with defective septa, is masked. Moreover, the negative effect on fitness of *pbp2x* knockdown can also be suppressed by knocking out PG synthesis genes *pbp1a, pbp1b* or *pbp2a* (Fig. 5A). PBP1a is involved in peripheral PG synthesis^38^, and we found it is synthetically lethal with *pbp2a*, indicating the two gene products may play a similar role, which was observed in a recent study^39^. Importantly, these data suggest that the rates of septal vs peripheral PG synthesis need to be balanced to ensure proper cell shape and division. However, if for instance the rate of septal PG synthesis is reduced, e.g., due to diminished PBP2x levels/activity, this can be rebalanced by slowing the rate of peripheral PG synthesis, indicating the existence of a delicate balance between septal and peripheral PG synthesis.

**Figure 5.**
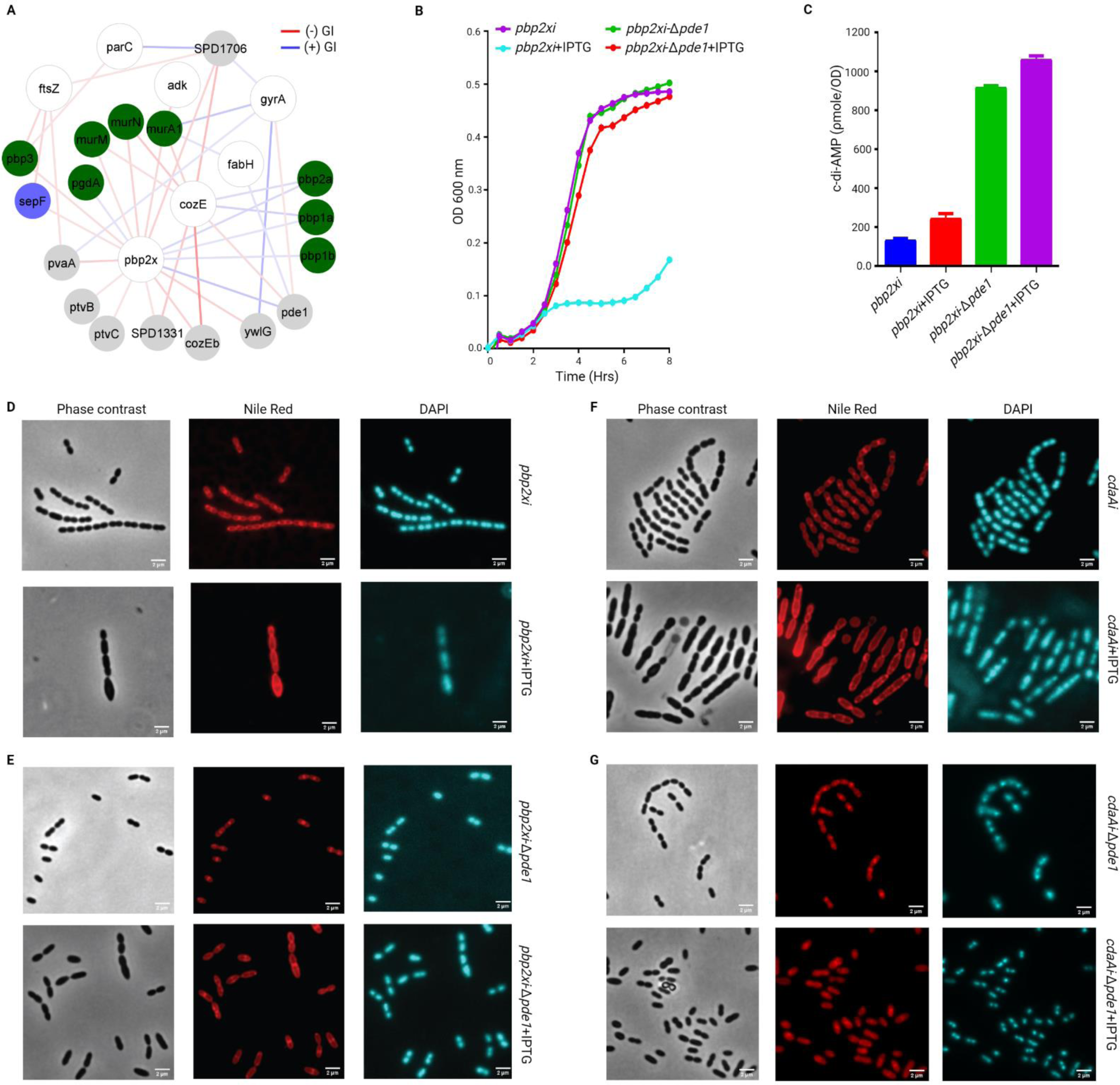
A *pde1* knockout mutant can buffer the negative phenotypic effects of *pbp2x* knockdown. **A.** A network highlighting the genetic interactions between *pbp2x* and nonessential genes involved in directly related processes including cell wall synthesis (green) and division (blue), as well as hypothetical genes (grey). Additional essential genes are highlighted that interact with the same nonessential genes as *pbp2x*. CRISPRi-targeted genes are presented as white nodes and nonessential genes as circles colored based on their bioprocess listed in Figure 2 with color code. Negative and positive genetic interactions (GI) are presented by red and blue edges, respectively. The intensity of edge color is proportional to the strength of the interaction z-score ranging from −10 to +10. **B.** Growth curves of *pbp2x*-CRISPRi (*pbp2xi)* and *pbp2xi*-Δ*pde1* without (violet and green) and with (cyan and red) 20 µM IPTG. Average of 3 replicate growth curves is plotted with standard error. **C.** c-di-AMP levels in *pbp2xi* and *pbp2xi*-Δ*pde1*, with and without 20 µM IPTG. **D.** Fluorescence microscopy images of *pbp2xi* with and without 100 µM IPTG. Representative phase contrast, Nile red (red), and DAPI (blue) stained micrographs are presented with 2 µm scale bar. **E.** Fluorescence microscopy images of *pbp2xi*-Δ*pde1* with and without 100 µM IPTG. **F.** Fluorescence microscopy images of *cdaAi* with and without 40 µM IPTG. **G.** Fluorescence microscopy images of *cdaAi*-Δ*pde1* with and without 40 µM IPTG.

The ability to control and rebalance PG synthesis is critically important for instance during cell division at which point synthesis needs to be refocused towards the septum. Our data shows that this rebalancing is mediated through *pbp3* (Penicillin-binding protein 3) which has a strong negative genetic interaction with *pbp2x* (Fig. 5A). *pbp3* is a class C PBP, it hydrolyzes D-Ala from the pentapeptide and delocalizes from the future division site thereby spatially directing availability of PG precursors towards the septum^40^. In the absence of PBP3, active PG precursors remain spread out across the membrane, reducing the effective concentration at the septum, while inhibiting *pbp2x* further disturbs this imbalance and exacerbates the problem. Furthermore, the importance of balanced PG-synthesis and the connection with cell division is highlighted by the strong negative interactions between *pbp2x* with *sepF* and *pvaA,* and *ftsZ* with *pbp3, sepF*, and *pvaA* (Fig. 5A). SepF has been shown to anchor and activate Z-ring assembly in *Bacillus subtilis*^41^, while *pvaA*, a lysozyme like protein, is important for proper localization of FtsZ^42^. The interactions with *sepF* confirm it has a similar function of activation of Z-ring assembly in *S. pneumoniae*. Overall, these strong negative interactions among PG synthesis genes and with Z-ring assembly related genes demonstrate that both Z-ring formation and localization, and septal and peripheral PG synthesis are intimately coordinated to successfully accomplish cell division.

### When cell wall synthesis is compromised, c-di-AMP can maintain growth and morphology by controlling turgor

CRISPRi-TnSeq shows that Δ*pde1* can suppress the effects of *pbp2x* knockdown, which we validated (Fig. 5B). *pde1* encodes a phosphodiesterase that hydrolyzes the second messenger cyclic di-AMP (c-di-AMP). We hypothesized that the absence of *pde1* increases intracellular c-di-AMP, which maintains growth and morphology under reduced levels of PBP2x. Indeed, both intracellular and extracellular c-di-AMP levels are high in Δ*pde1* (Fig. 5C, Supplemental Fig. 7). However, increased c-di-AMP levels in the growth medium alone are not sufficient to rescue *pbp2x* knockdown (Supplemental Fig. 8), suggesting that high intracellular c-di-AMP levels buffer against the inhibition of septal PG synthesis by decreasing intracellular turgor^43^. Fluorescence microscopy shows that *pbp2x* knockdown leads to enlarged cells at a coccus-to-rod transition state and with defective septa (Fig. 5D). Importantly, Δ*pde1* resolves this strong morphological change back to a wild type-like morphology (Fig. 5E), albeit with shorter chain-lengths, suggesting that chain-length determinants, such as LytB, are altered in Δ*pde1*^44^. c-di-AMP synthase or diadenylate cyclase (*cdaA*) synthesizes and maintains cellular c-di-AMP levels. We hypothesized that CRISPRi knockdown of *cdaA* reduces c-di-AMP levels and impacts cell morphology, and we should be able to rescue morphology by knocking out *pde1*. To test this, *cdaA* was knocked down in both wild-type and Δ*pde1* backgrounds. Indeed, the knockdown of *cdaA* inhibited the growth of wild-type but the impact of knockdown was mild in Δ*pde1* (Supplemental Fig. 9). Enlarged cells with coccus-to-rod transition, uneven chromosome distribution, and defective septa were observed when *cdaA* was knocked down in the wild-type background (Fig. 5F). But knockdown of *cdaA* did not impact the morphology of Δ*pde1* (Fig. 5G). The impacts of *cdaA* knockdown were thereby to some extent similar to what was observed when *pbp2x* was knocked down.

As a secondary messenger, c-di-AMP regulates multiple biological processes^45–47^. However, its role in cell wall synthesis has not been explored in *S. pneumoniae*. It has been shown that intracellular c-di-AMP directly interacts with effector protein CabP, which reduces the activity of the TrkH potassium transporter, thereby inhibiting the uptake of K^+^ ^48^ and affecting cellular turgor^43^. We hypothesized that the increased c-di-AMP levels of Δ*pde1* lead to reduced intracellular K^+^, which in turn reduces cellular turgor and thereby rescues growth and morphology when *pbp2x* is knocked down. Indeed, when the extracellular K^+^ concentration is increased, the effects of Δ*pde1* on cellular turgor, and the rescuing effect on *pbp2x* knockdown is nullified (Fig. 6A). Moreover, growth in the presence of valinomycin, a K^+^-carrier^49^ that promotes K^+^ uptake independent of a TrkH transport system also neutralizes the rescuing effect of Δ*pde1* on *pbp2x* knock down (Fig. 6B). This indicates that a lower intracellular K^+^ concentration, for instance mediated by increased levels of c-di-AMP, can protect cells from cell wall synthesis perturbation/inhibition (Fig. 6C). Importantly, these data show that intracellular c-di-AMP plays a protective role in *S. pneumoniae* by controlling turgor, which possibly works in a similar fashion in *Bacillus subtilis*, and *Listeria monocytogenes*^50^, and can thereby modulate the susceptibility to perturbation of the cell wall synthesis machinery, including by inhibitors such as penicillin.

**Figure 6.**
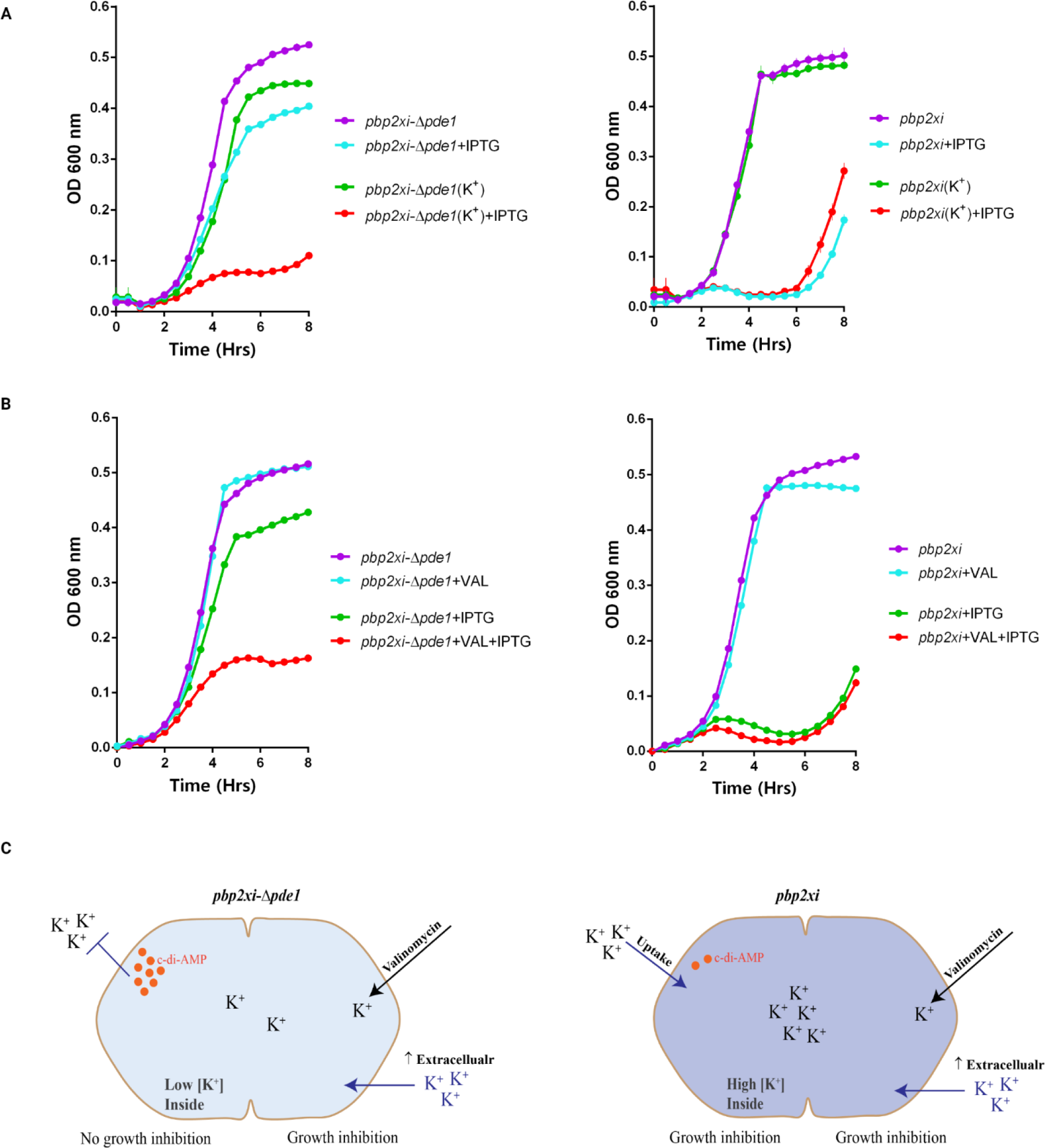
*pbp2x* knockdown can be rescued by decreasing K^+^ uptake and turgor. **A.** Growth curves of *pbp2xi* and *pbp2xi*-Δ*pde1* in the presence and absence of IPTG and/or additional potassium (K^+^). While Δ*pde1* can mask the negative fitness effect of *pbp2x* knockdown (see Figure 5B), the addition of K^+^ nullifies this masking effect. Average of 3 replicates are plotted with standard error. **B.** Growth curves of *pbp2xi* and *pbp2xi*-Δ*pde1* in the presence and absence of IPTG and/or valinomycin, a K^+^ -carrier. Average of 3 replicates are plotted with standard error. **C.** Modulation of the K^+^ concentration in *pbp2xi* and *pbp2xi*-Δ*pde1*. c-di-AMP reduces the activity of the TrkH potassium transporter, thereby inhibiting the uptake of K^+^ and affecting cellular turgor. Increased c-di-AMP levels in *pbp2xi*-Δ*pde1* in the presence of IPTG (i.e., *during pbp2x* knockdown) leads to reduced intracellular K^+^, which reduces cellular turgor and rescues growth and morphology.

### Midcell PG synthesis is modulated by CozEa and CozEb

As would be expected, in the interaction network *pbp2x* is part of a tight cluster of genes that are either directly or indirectly involved in cell wall metabolism and maintenance. *cozEa* is a gene that shares multiple interactions with *pbp2x*, which through ‘guilty by association’, indicates a shared mechanistic involvement. For instance, both have negative interactions with *murM*, *murN* and *murA* and suppressive interactions with *pbp1a* and *pbp2a* (Fig. 7A), which indicates that *cozEa* may also affect septal PG synthesis and thereby assist in rebalancing disturbances in peripheral PG synthesis. Moreover, CozEa was recently proposed as a member of the MreCD morphogenic complex of *S. pneumoniae*, controlling cell shape by positioning PBP1a in the midcell^51^. These relationships are all confirmed in the interaction network (Fig. 7A), which further supports CozEa’s membership to the MreCD complex and a role in cell morphology. Additionally, CozEa negatively interacts with *divIVA* (Fig. 7A), which is another gene involved in cell shape^52^. Furthermore, titration of *cozEa* makes *cozEb* extremely important (Fig. 7A), indicating (partial) functional redundancy, which is supported by a recent report showing that overexpression of CozEb can, to some extent, compensate for the lack of CozEa^53^. *cozEb* similarly negatively interacts with *pbp2x*, as knockdown of *pbp2x* arrests growth of Δ*cozEb* (Fig. 7B). We have previously shown and confirmed here (Supplemental Fig. 10 and 11), that Δ*cozEb* is hypersusceptible to daptomycin^17^, a cell wall disruptor. Additionally, it can be targeted *in vivo* with an antibody^4^, suggesting it is present in the cell wall and has a functional role in cell wall maintenance and/or synthesis. A green fluorescence protein (GFP)-tagged version of CozEb was constructed (Supplemental Fig. 12) to help visualize its localization. Time-lapse fluorescence microscopy in the presence and absence of a sub-inhibitory concentration of daptomycin suggests that early on during division CozEb can be mostly found around the septum before it distributes along the envelope of the cell (Fig. 7C). This patched distribution along the cell membrane was further confirmed with 3D-Structural Illumination Microscopy (3D-SIM) (Fig. 7D). To get more insight into the dynamics of CozEb overtime, the localization of GFP-CozEb was followed every 45 ms in live cells using Total Internal Reflection microscopy and Highly Inclined Laminated Optical sheet illumination (TIRF/HILO). This shows that CozEb is highly dynamic and seems to move rapidly along the membrane without following a clear pattern, but it also accumulates at the septum, where it seems to assemble/disassemble rapidly (Fig. 7E, Supplemental Movie 1). Importantly, the addition of daptomycin does not seem to influence CozEb dynamics. Recently, it was shown that both CozEa and CozEb are present in a complex with PBP1a and are possibly involved in maintaining midcell localization of PBP1a^53^. The dynamic nature of CozEb suggests a function of alternating interactions, possibly with a variety of proteins including PBP1a, PBP2x and CozEa. Moreover, the mobile nature and genetic interactions with multiple peripheral PBPs suggest that CozEa and CozEb may restrict class A PBPs in the midcell, possibly through physical interactions.

**Figure 7.**
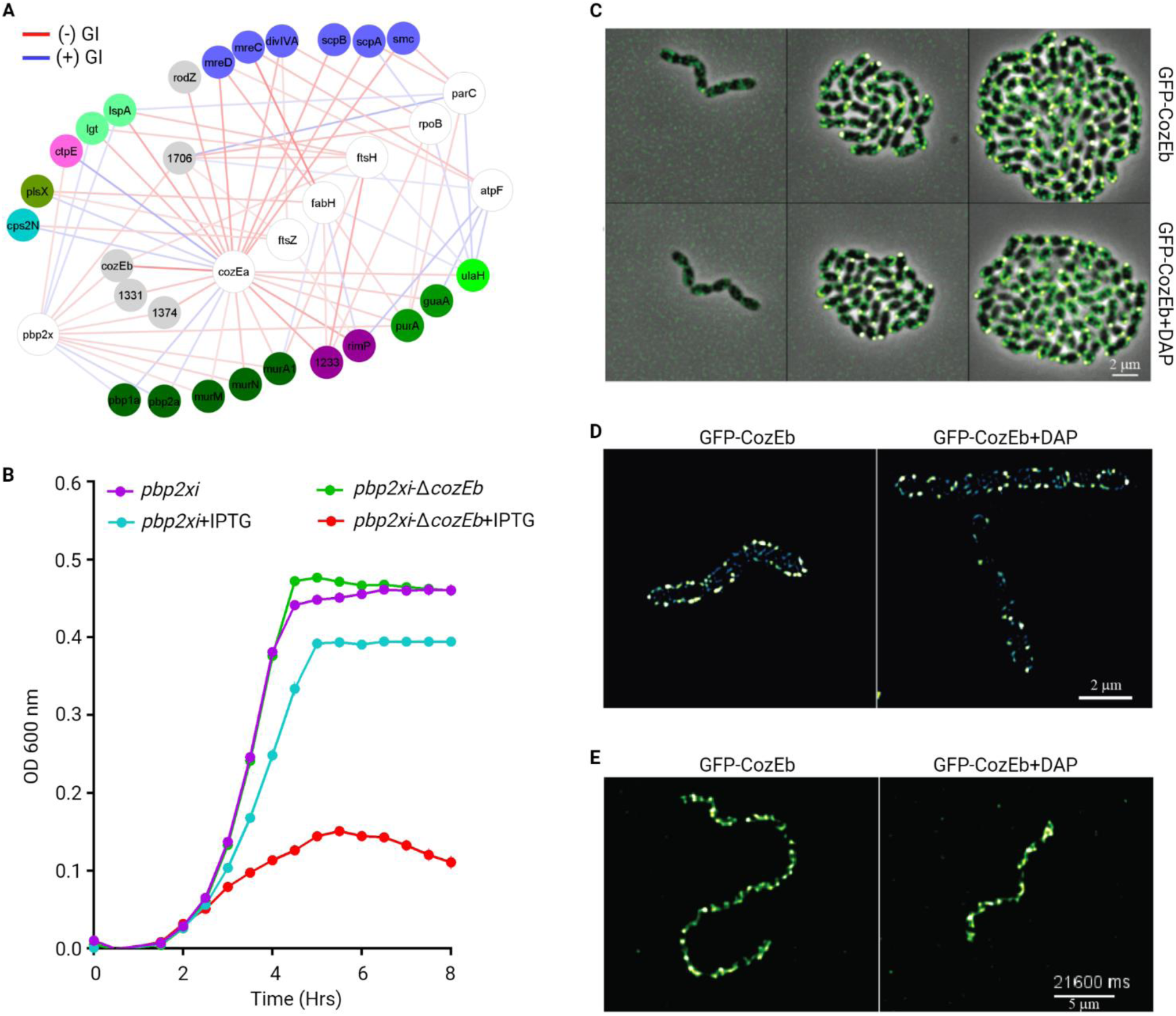
*cozEb* is a dynamic protein in the cell wall with a strong negative interaction with *pbp2x*. **A.** A network highlighting the genetic interactions between *cozEa* and nonessential genes involved in directly related processes including cell wall synthesis (green) and division (blue), as well as hypothetical genes (grey). Additional essential genes are highlighted that interact with the same nonessential genes as *cozEa*. CRISPRi-targeted genes are presented as white nodes and nonessential genes as circles colored based on their bioprocess listed in Figure 2 with color code. Negative and positive genetic interactions (GI) are presented by red and blue edges, respectively. The intensity of edge color is proportional to the strength of the interaction z-score ranging from −10 to +10. **B.** Growth curves of *pbp2xi* and *pbp2xi*-Δ*cozEb* in the presence and absence of IPTG. Average of 3 replicates are plotted with standard error. **C.** Time-lapse fluorescence microscopy of P_Zn_-*gfp*-*cozEb* in the absence and presence of 4 μg/mL daptomycin (DAP). Representative micrographs are presented with 2 µm scale bar. **D.** 3D-Structural illumination microscopy (3D-SIM) images of GFP-CozEb in live cells in the absence (left panel, chain of 3 cells) and presence (right panel, chain of 7 cells) of 5 μg/mL daptomycin. Fluorescent signal can be found around the cell. Signal intensity represented by colors, where blue is lower, green medium and white higher signal. Representative micrographs are presented with 2 µm scale bar. **E.** Snapshot from time-lapse of GFP-CozEb (green signal) in live cells (cells outlines not shown) at 37°C using TIRF (HILO) microscopy in the absence (left panel, chain of 16 cells) and presence (right panel, chain of 8 cells) of 5 μg/mL daptomycin. Images were taken every 45 ms. Representative micrographs are presented with 5 µm scale bar. Movie can be found as Supplemental Data Movie 1.

### Morphology and cell wall synthesis are maintained by coordinated actions of CozEa and RodZ

As highlighted in the interaction network, CozEa’s genetic interactions go beyond genes that are involved in PG synthesis and extends towards those involved in cell shape and morphology, including RodZ (Fig. 7A, 8A). In rod shaped species like *Escherichia coli*, RodZ regulates the localization of MreB in a concentration dependent manner^54^. MreB is a cytoskeleton actin-like protein that spatially synchronizes cell wall insertion directing the cell towards a rod shape^54^. The ovococcus *S. pneumoniae* lacks actin-like MreB, but does possess RodZ, which seems to have an important role in cell wall synthesis^55, 56^, and due to the lack of MreB it could have additional and/or other roles in ovococci. While a RodZ knockout does not result in a significant growth defect (Fig. 8A), it does trigger morphological changes, including irregularities in chain length, cell size, and chromosome distribution (Fig. 8B, C). A *rodZ* knockout combined with a *cozEa* knockdown, results in drastically reduced growth, and exacerbated morphological defects, with reduced chain lengths, ballooning, and lysis (Fig. 8A, B, C). Furthermore, Δ*rodZ* is highly sensitive to penicillin exposure (Supplemental Table 7), and *rodZ* shares multiple interactions with *mreCD* (identified here and other studies), including with *pbp1a*, MltG, EloR and *cozEa* (Fig. 8D)^55, 56^. As described above, MreCD is a morphogenic complex and like CozEa it is implicated in peripheral PG synthesis^51, 57^. This suggests that there is a functional link between *cozEa*, *rodZ*, and *mreCD,* which are all key to maintaining proper morphology, and that *rodZ* may share some redundancy with *cozEa* and possibly with *mreCD*.

**Figure 8.**
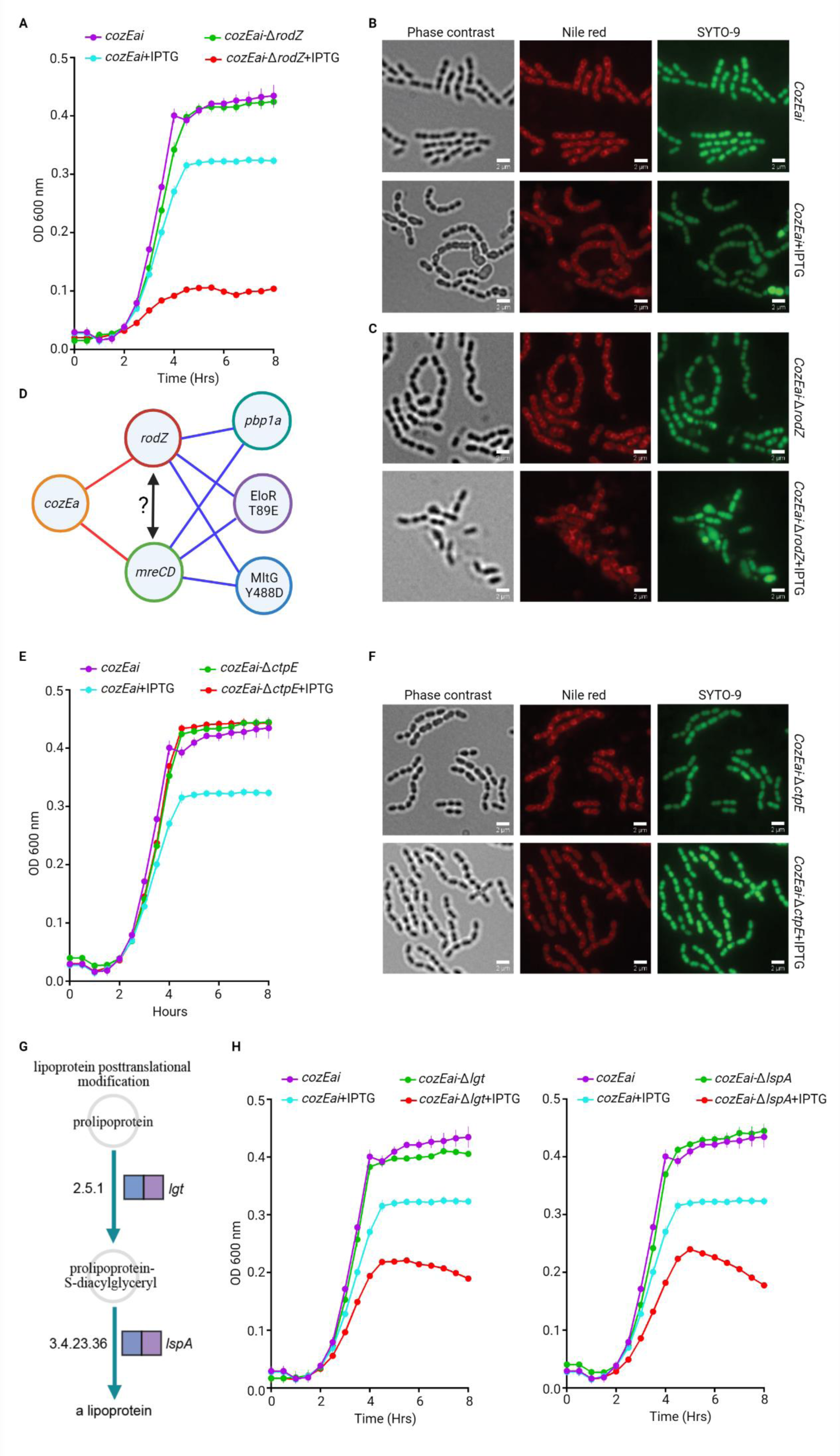
*cozEa* interacts with different genes involved in maintaining cell wall integrity. **A.** Growth curves of *cozEai* and *cozEai*-Δ*rodZ* in the presence and absence of IPTG. Average of 3 replicates are plotted with standard error. **B.** Fluorescence microscopy images of *cozEai* with and without IPTG. Representative phase contrast, Nile red (red), and SYTO-9 (green) stained micrographs are presented with 2 µm scale bar. **C.** Morphological profiles of *cozEai*-Δ*rodZ* with and without 100 µM IPTG. **D.** Summary of genetic interaction profiles of *rodZ* and *mreCD*. Negative and positive genetic interactions are presented by red and blue edges, respectively. **E.** Growth curves of *cozEai* and *cozEai*-Δ*ctpE* in the presence and absence of IPTG. Average of 3 replicates are plotted with standard error. **F.** Morphological profiles of *cozEai* and *cozEai*-Δ*ctpE* with and without 100 µM IPTG. **G.** Lipoprotein posttranslational modification pathway of *S. pneumoniae,* including nonessential genes *lgt* and *lspA*. Both genes were identified in CRISPRi-TnSeq to have a negative interaction with *cozEa*. **H.** Growth curves of *cozEai, cozEai*-Δ*lgt* and *cozEai*-Δ *lspA* in the presence and absence of IPTG. Average of 3 replicates are plotted with standard error.

### CozEa controls cell wall integrity through CtpE and calcium homeostasis

With the strong positive interaction between *cozEa* and *ctpE* (Fig. 7A, 8E), *cozEa*’s role seems to further extend into maintaining envelope integrity by affecting calcium homeostasis. In *Mycobacterium smegmatis*, *ctpE* is key in Ca^2+^ uptake and homeostasis during limited Ca^2+^ availability and is possibly involved in maintenance of envelope integrity due to its genome localization in an operon with genes predicted to be involved in membrane and cell wall biosynthesis and integrity^58^. While knockdown of *cozEa* has a mild impact on morphology and growth, these defects can be completely suppressed by a *ctpE* knockout (Fig. 8E, F). This suggests that a reduced amount of CozEa results in CtpE dysregulation and changes in intracellular calcium that lead to defects in cell envelope integrity. Importantly, under conditions where calcium concentrations in the environment are high, disabling *ctpE* seems to restore calcium homeostasis and compensates the negative fitness effect of CozEa knockdown, suggesting that CozEa has a direct or indirect regulatory effect on CtpE. Moreover, negative interactions exist between *cozEa* and at least two lipoprotein maturation related genes, *lspA* and *lgt*^59, 60^(Fig. 7A; 8G, H), which further confirms a role in maintaining envelope integrity.

### Chromosome segregation promotes DNA decatenation and modulates cell morphology

Finally, the tight integration of seemingly disparate processes are further underlined by negative interactions between *cozEa* and *rimP* and SPD1233, two translation related genes, as well as with a cluster of DNA replication associated genes, including *smc*, *scpA*, *parC*, *parB* and *topA*. While these connections underscore the relationship between the transfer, replication and translation of genetic information, cell wall synthesis, integrity and cell division, the tight cluster of DNA replication associated genes reveals some important relationships between these genes. For successful cell division, the decatenated chromosome needs to be segregated to daughter cells. Previously, *smc* and *parB,* have been shown to influence chromosome segregation in *S. pneumoniae*^36^, while ParC is involved in chromosome decatenation^61^. Here, we identify and validate a negative interaction between *smc* and *parC* (Fig. 9A, B). Fluorescence microscopy shows that *parC* titration results in anucleate cells and uneven chromosome distribution (Fig. 9D), highlighting that chromosome distribution is dependent on decatenation by *parC*. Deletion of *smc* results in enhanced chaining (Fig. 9E), indicating that SMC-mediated chromosome segregation affects downstream cell division. Knockdown of *parC* combined with Δ*smc,* produces visual deleterious effects, that combines alterations in chromosome distribution and cell morphology (Fig. 9E). Additionally, the negative interaction between *parC* and the partitioning protein ParB (Fig. 9A, C, F), whose main role is to recruit SMC to the *oriC*, confirms functional dependency between chromosome decatenation and segregation, and the critical link with cell division, and cell wall synthesis.

**Figure 9.**
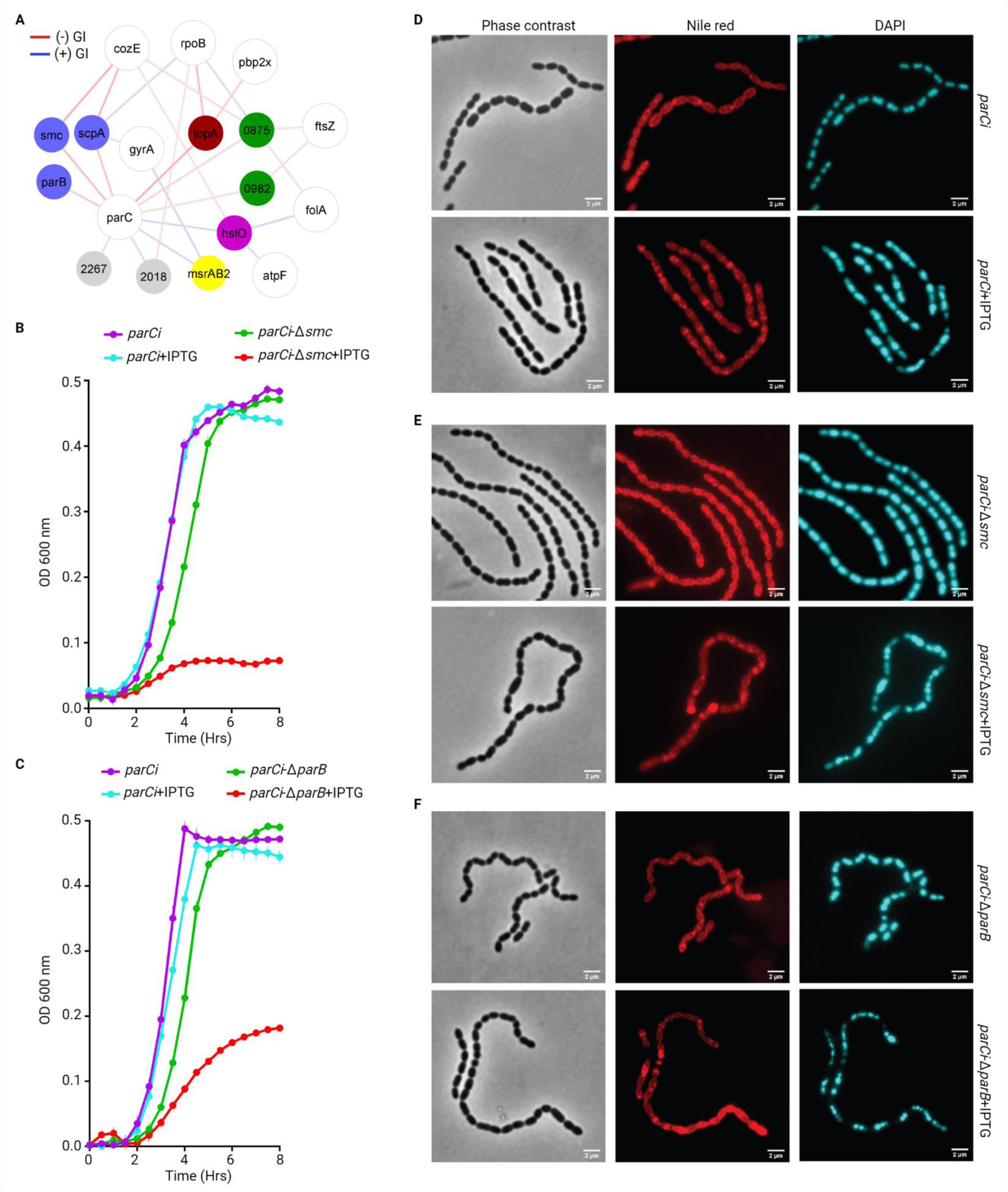
CRISPRi-TnSeq analysis of *parC*-CRISPRi strain. **A.** A network highlighting the genetic interactions between *parC* and non-essential genes involved in cell wall synthesis (green), division (blue), as well as hypothetical genes (grey). Additional essential genes are highlighted that interact with the same non-essential genes as *cozEa*. CRISPRi-targeted genes are presented as white nodes and nonessential genes as circles colored based on their bioprocess listed in Figure 2 with color code. Negative and positive genetic interactions (GI) are presented by red and blue edges, respectively. The intensity of edge color is proportional to the strength of the interaction z-score ranging from −10 to +10. **B.** Growth curves of *parCi* and *parCi*-Δ*smc* in the presence and absence of IPTG. Average of 3 replicates are plotted with standard error. **C.** Growth curves of *parCi* and *parCi*-Δ*parB* in the presence and absence of IPTG. Average of 3 replicates are plotted with standard error. Fluorescence microscopy images of of *parCi* (**D**) *parCi*-Δ*smc* (**E**) and *parCi*-Δ*parB* (**F**) with and without IPTG. Representative phase contrast, Nile red (red), and DAPI (blue) stained micrographs are presented with 2 µm scale bar.

## Discussion

By exploiting the combined power and flexibility of essential gene knockdown and nonessential gene knockout, we develop CRISPRi-TnSeq to map bacterial essential-nonessential genetic interaction networks. By focusing on key biological processes including metabolism, DNA replication, transcription, cell division and cell envelope synthesis we screen over 24,000 gene-gene pairs, which leads to the identification of 1,334 significant genetic interactions. Network analyses show that essential gene knockdown leads to genetic interactions with functionally linked genes, indicating that genetic interaction data can be leveraged to verify known associations between genes as well as identify new ones. To establish the utility of CRISPRi-TnSeq we mine the network and explore and validate a variety of new interactions. First, by ranking the nonessential genes based on their number of interactions, 17 nonessential genes emerge that interact with more than half of the targeted essential genes, which makes them the most pleotropic in the dataset. In other species genes with pleiotropic functions have been shown to work as genetic capacitors that can buffer against stressors^8^. Here, these genes also seem to introduce robustness into the system by dampening the phenotypic outcome during genetic and environmental perturbations. Consequently, such genes can make an entire system more vulnerable, including by increasing sensitivity to antimicrobials, and thus they could be valuable as targets for combination-drugs. Furthermore, by focusing our analyses on interactions involved in cell wall synthesis, cell division and DNA replication multiple key biological insights are uncovered. For instance, we show that knockdown of an essential gene product can be compensated through nonessential gene-redundancy and flux rebalancing to make the pathway less dependent on the essential gene. This balancing act is further underscored by PG synthesis, for which we show that when septal PG synthesis is reduced due to the knockdown of PBP2x overall fitness decreases, which can be further aggravated by precursor gene knockouts such as *murM*. In contrast, fitness can be rescued by simultaneously slowing down peripheral PG synthesis by knocking out genes such as *pbp1a*, highlighting the importance of balanced and fine-tuned processes. These data are in line with a recent study in which we showed how gene-essentiality can depend on the genetic background and how essential genes and processes can evolve to become nonessential through genetic rewiring that can in certain cases be achieved by even a single or several SNPs and/or relatively small changes in expression^16^. Moreover, CRISPRi-TnSeq underscores the interdependencies that exist genome-wide between distinct processes. While cell division is a complex process on itself, the genetic interaction network highlights the dependency on coordinated action between multiple processes including cell wall synthesis and organization (e.g., *pbp2x*, *pbp1a*, *murM*, *murN*, *cozEa*, *cozEb*), Z-ring assembly and localization (e.g., SepF, FtsZ) cell shape and morphology (e.g., RodZ) and chromosome decatenation and segregation (e.g., *smc*, *scpA*, *parC*, *parB*). Importantly, interactions with, *pde1*, *cdaA*, and *ctpE*, show how biometals such as calcium and potassium and the signaling molecule c-di-AMP, can (in)directly control these processes through for instance turgor-control, thereby ensuring survival and bacterial growth, albeit at a slower rate. Importantly, a similar role of c-di-AMP has been reported in *Bacillus subtilis*, and *Listeria monocytogenes*^50^, indicating a conserved protective function that fine-tunes and balances a variety of processes that contribute to cell growth and division.

In conjunction with other tools, CRISPRi-TnSeq highlights the importance of mapping genetic interactions. On one hand the identification of suppressor interactions is a strength of CRISPRi-TnSeq as it uncovers hidden redundancy in pathways and/or processes. This, in turn, can be an indicator of the ease in which an organism can find a solution (e.g., the likelihood resistance may emerge) against a drug targeting that pathway. In contrast, aggravating interactions, whether they are within or between pathways or processes, highlight the dependencies within a genome and thereby potential drug targets that would work synergistically. Importantly, while CRISPRi-TnSeq is a powerful new addition to the geno-phenotyping genomics toolbox it will by no means be the last. For instance, we envision two alternative and upgraded versions: 1. By creating double knockout Tn-libraries, genetic interaction networks could be generated on a genome wide scale and in high-throughput. However, standard Tn-Seq methods cannot determine which two genes are knocked-out in the same background, and thus the Tn-context of a double knockout library would be lost; 2. By incorporating two guide RNAs into a single cell, a dual-guide CRISPRi tool would have the potential to become a broadly applicable approach for genome-wide genetic interaction mapping. We are actively working on both approaches as we foresee that each will have its own advantages and disadvantages with respect to questions that can be answered as well as their differential ease of implementation and optimization into a new strain or species. We expect the implementation of CRISPRi-TnSeq to be relatively easy for many bacteria, i.e., if both tools are implemented in a species, then, as we show here, they merely need to be combined. Moreover, the experimental set-up and execution, sample prep, data collection and analyses we outlined here can be used as a detailed guideline without the need for much further development. We thus believe that CRISPRi-TnSeq will prove to be a powerful tool that can untangle fundamental network relationships between essential and nonessential parts of the genome and thereby uncover new biological insights.

## Materials and Methods

### Strains, growth conditions, and transformation

CRISPRi strains of *Streptococcus pneumoniae* D39V (NZ_CP027540.1) were adopted from a previous study^26^ and are detailed in Supplemental Table 1. *Streptococcus pneumoniae* D39V and its derivatives were cultivated either in C+Y media, pH 6.8^29^, Todd Hewitt broth supplemented with yeast extract (THY), 5 µg/mL of Oxyrase (Oxyrase Inc.) and 150 U/mL Catalase (Worthington Bio Crop), or on 5% Sheep’s blood agar plates at 37^0^C in a 5% CO_2_ atmosphere. Unless otherwise indicated, for growth studies, cells were grown up to the mid exponential phase, diluted in the same growth media, and incubated in a BioSpa automated incubator with integrated plate reader (BioTek). Growth curves were plotted using GraphPad Prism9. Transformation of *S. pneumoniae* was performed as previously described^14^.

### Transposon mutant library construction and selection experiments

Transposon mutant libraries of individual CRISPRi strains using the mariner transposon Magellan6 were constructed as previously described^3, 13, 14^. The transposon carries a kanamycin resistance cassette that lacks a transcriptional terminator therefore allowing for readthrough transcription and has stop codons in all three frames in either orientation to prevent aberrant translation products. DNA was isolated from individual CRISPRi strains followed by *in vitro* transposition. Transposed DNA was used to transform the corresponding CRISPRi strain leading to the construction of a mutant library. Six independent transposon mutant libraries, each consisting of ∼10,000 mutants, were constructed for each CRISPRi strain. Mutant libraries were grown in C+Y media up to mid-exponential phase, which represents the initial population (T1) in a Tn-Seq experiment. Subsequently, a culture was diluted to OD 0.003 and grown in the same media in the absence and presence of two sub-inhibitory concentrations of IPTG or different antibiotics. Cells were harvested after 6-8 doublings, when culture density reached ∼0.8 OD.

### Expression analysis

Total RNA was isolated from cells grown with and without IPTG using the RNeasy mini kit (Qiagen. Inc), followed by DNase treatment with the TURBO DNA-*free* kit (Invitrogen). cDNA was synthesized from 1 µg RNA with random hexamers using the iScript Reverse Transcription Supermix (Bio-Rad). Quantitative PCR was performed on a MyiQ (Bio-Rad) using iTaq Universal SYBR Green Supermix (Bio-Rad). Each sample was measured in technical and biological replicates, and CRISPRi target gene expression was normalized against the 50S ribosomal RNA gene SPD2031 (*rplI*).

### Analyses of CRISPRi-transposon mutant libraries

Mutant libraries for each CRISPRi strain were grown in the absence and presence of IPTG followed by DNA isolation using the DNeasy Blood and Tissue Kit (Qiagen, Inc). Tn-Seq sample preparation, Illumina sequencing and fitness calculations were performed as described previously^3, 13, 14^. Importantly, fitness for each insertion mutant (*W_i_*) is calculated so that it represents the actual growth rate per generation making fitness independent of time, which allows comparisons between conditions and strains^3, 13^. To determine whether fitness values are significantly different between conditions, three conditions are imposed: 1. *W_i_* is calculated from 3 or more data points; 2. The fitness difference between conditions is larger than 0.1 (10%), and 3. the fitness difference is statistically significant in one sample *t*-test with Bonferroni or Benjamini-Hochberg correction for multiple testing. All fitness change values were presented in volcano plot using Vega-Lite^62^.

### Genetic interaction mapping

Genetic interactions of two genes are defined as a deviation from the multiplicative model. If individual knockout mutants of genes *a* and *b* have fitness (*W_a_*) and (*W_b_*), respectively, a genetic interaction is scored when fitness of the double mutant (*W_ab_*) deviates from the multiplicative fitness, (*W_a_* x *W_b_*)^11, 12^. In CRISPRi-TnSeq, the fitness difference between the absence (*W_noIPTG_*) and presence (*W_IPTG_*) of IPTG are calculated for the transposon mutant of individual nonessential genes. Since CRISPRi knockdown of an essential gene target reduces the growth of the entire mutant library, the relative fitness of individual insertion mutants remains similar in the presence of IPTG unless the corresponding essential and nonessential gene pair genetically interact. *W_ab_* should thus be significantly different from *W_a_* x *W_b_*, and Δ*w (*Δ*w = W_IPT G_* - *W_noIPTG_*) represents the size of the fitness effect. The impact on growth due to the essential gene knockdown depends on many factors, including stability and abundance of the corresponding gene product. To make genetic interaction values comparable between different datasets produced for individual CRISPRi targets, individual fitness difference (Δ*w*) and genetic interaction datasets were z-score normalized. All significant genetic interactions with p-adj ≤ 0.05 were mapped in a genetic interaction network using Cytoscape^63^. It should be noted that random transposon insertions are likely in the large *dcas9* gene. Those mutants are expected to be identified with increased fitness in CRISPRi-TnSeq due to the lack of an active knockdown system. As expected, *dcas9* mutant fitness increased with increased IPTG concentrations for each library (Supplemental Table 3), which confirms successful target knockdown and growth inhibition of the entire mutant library except the mutants of *dcas9* gene. Therefore, *dcas9* essentially serves as a positive control in the CRISPRi-TnSeq datasets.

### Genetic interaction distributions, enrichment, and visualization

*S. pneumoniae* D39 genome was plotted using Circos 0.69. The distribution of interactions across the genome was determined for each 1 kb region and plotted as a histogram. Enrichment was determined for each 50kb region in the genome by comparing it to the distribution of interactions of random regions within the genome.

The Jaccard index is determined, which is defined as the size of the intersection of the significant gene pool divided by the size of the union of the significant gene pool. Gene set enrichment analysis (GSEA)^64^ was performed to test the overrepresentation of genes from specific biological process(es) and were summarized in a heatmap as q-values indicating false discovery rates. Pathway analyses were performed for individual datasets using BioCyc^65^. Genetic and chemical genetic interaction datasets were visualized with Cytoscape^63^.

### Gene deletion mutant construction

Gene knockout mutants were constructed by substituting the entire coding sequence with a chloramphenicol resistance cassette constructed through overlap extension PCR as previously described^13, 66^. Deletion mutant strains and oligos can be found in Supplemental Tables 1 and 9, respectively.

### Construction of *gfp*-labeled strains

Strains and primers used in this study are listed in Supplemental Tables 1 and 9, respectively. Strain CG89 (*S. pneumoniae* TIGR4, *bgaA*::P_Zn_-*m(sf)gfp*-*sp_1505*) was constructed by transformation of *S. pneumoniae* TIGR4 wild type with plasmid pCG15. This plasmid was built by ligating the product amplified by PCR with primers SP1505_F-SpeI and SP1505-R-NotI from *S. pneumoniae* DNA, into plasmid pCG6, allowing for N-terminal GFP fusion of SP1505 under the control of a ZnCl_2_-inducible promoter (P_Zn_). Strain CG90 (TIGR4, *sp1505*::*kan*-*m(sf)gfp-sp1505*) was built by transformation of TIGR4 wild type with product *kan-m(sf)gfp-sp1505* in order to replace the original SP_1505 gene. The DNA product was obtained by Gibson assembly of three PCR fragments: (1) *m(sf)gfp-sp1505* amplified with primers SP_1505-up-Htra-GFP-F and SP_1505-down-R on CG89 genomic DNA; (2) *kan*, kanamycin resistance marker, amplified with primers kan-F and kan-R-SP_1505-up and (3) *sp1505* upstream region amplified with primers SP_1506-down-F and SPD_1506-up-R-kan on TIGR4 wild type genomic DNA.

### Cyclic-di-AMP assay

Strains were grown in C+Y media at 37°C with 5% CO_2_ up to OD_600nm_ 0.2-0.4. Cultures were diluted to 0.003 OD in C+Y and regrown with and without IPTG up to 0.5-1.0 OD. 5 OD x mL volume of culture were pelleted by centrifugation, cell pellets were resuspended in lysis buffer (20 mM Tris pH 7.4, 1% Triton X-100, 100 µg/mL lysozyme, 5 U/mL DNase I) followed by incubation at room temperature for 20 minutes. Lysates were centrifuged at 15,000 x g for 5 minutes at 4°C. Supernatants were used in a cyclic-di-AMP ELISA assay using kit (Cayman chemicals, catalog number 501960) and following manufacturer instructions. This kit measures cellular cyclic-di-AMP against HRP-tagged cyclic-di-AMP by competitive ELISA binding. Cyclic-di-AMP levels were calculated based on the standard curve, normalized by culture OD and presented in ρmole/OD.

### Fluorescence microscopy of CRISPRi strains and mutants

Morphological changes associated with the knockdown of target genes were analyzed by microscopy as described previously^26^. Briefly, cells were grown in C+Y at 37°C for 2.5 to 3 hours with and without IPTG. 1 µL of 1 mg/mL Nile red (membrane dye) was added to 1 mL of cell suspension and incubated at room temperature for 4 min, followed by 1 µL of 1 mg/mL of DAPI (DNA dye) and incubation for another minute. Next cells were precipitated by centrifuging at 8,000 x g for 2 min and re-suspended in fresh 30 µL C+Y medium. 0.5 µL of cell suspension was spotted on a PBS agarose pad prepared on the slide. Finally, cells were visualized with a fluorescence microscope Olympus IX83 (Olympus) or Deltavision Elite (GE Healthcare). Microscopy images were analyzed with Image J^67^.

### Time-lapse microscopy

*S. pneumoniae* TIGR4 strains were pre-grown in C+Y medium at 37°C until OD_600nm_ = 0.3. Cells were diluted 100-fold in fresh C+Y medium, supplemented when appropriate with daptomycin, and cultured until OD_600nm_ = 0.1. Cells were harvested by centrifugation for 1 min at 10,000 x g, washed once in fresh C+Y medium, and re-suspended into 1/10^th^ of the centrifuged volume. 1 μL was spotted onto a C+Y-10% polyacrylamide pad (incubated twice for 1h in C+Y medium supplemented with daptomycin when applicable) inside a Gene Frame (Thermo Fisher Scientific) sealed with a cover glass. The resulting slide was then placed into the microscope chamber pre-incubated at 30°C or 37°C. Phase contrast time-lapse microscopy, was performed as described before^68^ using a DV Elite microscope (GE Healthcare) with a sCMOS (PCO-edge) camera and a DV Trulight solid-state illumination module (GE Healthcare), and a ×100/1.40 oil-immersion objective. Conventional epifluorescence time-lapse microscopy was performed on the same microscope using a GFP filter (Ex: 475/28 nm, BS: 525/80 nm, Em: 523/36 nm). TIRF (HILO) fluorescence time-lapse microscopy was performed on a Leica DMi8 microscope with a sCMOS DFC9000 (Leica) camera using a 100x oil-immersion Plan APO TIRF objective, a 488 nm excitation laser module and a GFP filter (Ex: 470/40 nm Chroma ET470/40x, BS: LP 498 Leica 11536022, Em: 520/40 nm Chroma ET520/40 m). Images were acquired with either LasX v.3.4.2.18368 (Leica) or SoftWoRx v.7.0.0 (GE Healthcare) and processed with FIJI^69^.

### 3D-Structural illumination microscopy

Live bacterial cells for 3D-SIM were spotted onto PBS–10% acrylamide pads. Acquisition was performed using a DeltaVision OMX SR microscope (GE Healthcare) equipped with a ×60/1.42 NA objective (Olympus) and 488 nm excitation laser. Nine Z-sections of 0.135 μm each were acquired in Structure Illumination mode with 20 ms exposure and 25% laser power. The 135 images obtained were reconstructed with a Wiener constant of 0.01 using SoftWoRx (GE Healthcare).

## Data availability

All sequence data can be found in the NCBI Sequence Read Archive under the BioProject: PRJNA813307. Source data for the figures are available in the supplemental tables.

## Acknowledgement

DNA sequencing was performed at the Boston College Sequencing Core. We thank Jon Anthony for establishing the *Aerobio* sequencing analyses platform, and Federico Rosconi, Juance Ortiz-Marquez and Kristin Baker for valuable discussions. This work is supported by NIH R03 AI135737, R21 AI156203, U01 AI124302 and U19 AI158076 to T.v.O. Xue Liu is supported by the National Nature Science Foundation of China (NSFC, 82270012) and the Science and Technology Project of Shenzhen (JCYJ20220818095602006).

## Author contributions

B.J. and T.v.O designed the research and wrote the manuscript. B.J., X.L., J.D., H.P., B.L. and C.G. performed experiments and collected data. B.J. and D.L. performed all bioinformatic and statistical analyses. T.v.O. and J.W.V. supervised the project. All authors contributed to manuscript writing and approved the final paper.

## Competing interests

The authors declare no competing interests.

## Notes

### Competing Interest Statement

The authors have declared no competing interest.

